# Antagonistic interactions subdue inter-species green-beard cooperation in bacteria

**DOI:** 10.1101/2020.02.25.965665

**Authors:** Santosh Sathe, Rolf Kümmerli

## Abstract

Cooperation can be favored through the green-beard mechanism, where a set of linked genes encodes both a cooperative trait and a phenotypic marker (green beard), which allows carriers of the trait to selectively direct cooperative acts to other carriers. In theory, the green-beard mechanism should favor cooperation even when interacting partners are totally unrelated at the genome level. Here, we explore such an extreme green-beard scenario between two unrelated bacterial species – *Pseudomonas aeruginosa* and *Burkholderia cenocepacia*, which share a cooperative locus encoding the public good pyochelin (a siderophore) and its cognate receptor (green beard) required for iron-pyochelin uptake. We show that pyochelin, when provided in cell-free supernatants, can be mutually exchanged between species and provide fitness benefits under iron limitation. However, in co-culture we observed that these cooperative benefits vanished and communities were dominated by *P. aeruginosa*, regardless of strain background and species starting frequencies. Our results further suggest that *P. aeruginosa* engages in interference competition to suppress *B. cenocepacia*, indicating that inter-species conflict arising from dissimilarities at the genome level overrule the aligned cooperative interests at the pyochelin locus. Thus, green-beard cooperation is subdued by competition, indicating that inter-specific siderophore cooperation is difficult to evolve and to be maintained.

## Introduction

Explaining the origin and maintenance of cooperation is a challenge for evolutionary biology (Sachs et al. 2004; Nowak 2006; West et al. 2007; Leigh Jr 2010; Bourke 2011; Raihani et al. 2012). The puzzle is why should individuals carry out behaviors that bear some immediate costs to themselves but benefit others? And once established, how can cooperation be maintained given that non-cooperators (cheaters) may arise and exploit cooperators? Despite these challenges, cooperative behaviours are widespread and are observed at all levels of biological organizations (Frank 2007). For example, genes cooperate in genomes, cells cooperate in multicellular organisms and multicellular organisms come together to form cooperating societies as in the social insects and humans (Maynard Smith and Szathmary 1995; Bourke 2011; West et al. 2015).

Inclusive fitness theory suggests that cooperation can evolve if individuals, sharing the genes responsible for cooperation, interact with each other more often than expected by chance (Hamilton 1964; Gardner et al. 2011). This form of high genetic relatedness at the cooperating locus can either arise when the interacting individuals share recent common ancestry (leading to high genetic relatedness throughout the entire genome) or through the so-called green-beard effect, where the interacting individuals share the cooperative locus, but are unrelated at other parts of the genome (Dawkins 1976; Gardner and West 2010; West and Gardner 2013). The green-beard locus typically consists of one or a set of tightly linked genes with the following three properties: (a) they encode a cooperative behaviour and a detectable phenotypic marker (green-beard); (b) they enable the bearers to use the phenotypic marker to discriminate non-carriers from carriers displaying the same phenotypic marker; and (c) they allow green-beard carriers to selectively direct cooperative acts to other carriers (Gardner and West 2010; West and Gardner 2010; Gardner 2019).

The green-beard effect has attracted the attention of evolutionary biologists for many decades (Dawkins 1976; Ridley and Grafen 1981; Dawkins 1982; Rushton et al. 1984; Keller and Ross 1998; Jansen and van Baalen 2006; Sinervo et al. 2006; Biernaskie et al. 2011; Nonacs 2011; Biernaskie et al. 2013; Joshi and Guttal 2018; Madgwick et al. 2019), although green-beard loci were initially predicted to be rare for various reasons (Gardner and West 2010). First, it was hard to conceive how a gene or a set of tightly linked genes can simultaneously code for all the above-mentioned properties. Second, the linkage between the green-beard genes can break and therefore cheaters (i.e. individuals with a false beard who only express the phenotypic marker but do not cooperate) can evolve and spread in the population. Third, as the green-beard effect excludes non-carriers from the benefits of cooperation it is thought that green-beard loci provide large fitness benefits and should thus quickly fix in populations and thereby lose their discriminatory function. Fourth, the green-beard effect can lead to conflicts between the green-beard locus (high relatedness) and the rest of the genome (low relatedness), as the evolutionary interests of the two parts are not necessarily aligned. These four reasons combined probably explain why only a limited number of green-beard genes have been discovered so far. One example is the *GP*9 locus in the red fire ant *Solenopsis invicta* (Keller and Ross 1998), where workers that carry the *b* allele discriminate against and kill homozygous *BB* queens, whereas *Bb* queens remain unharmed. In this case, the *b* allele is a harming green-beard that specifically targets non-carriers, but cannot get fixed in populations because it is lethal in homozygous individuals (Keller and Ross 1998). Other green beards are the *csA* and *tgr* loci in *Dictyostelium discoideum* (Queller et al. 2003; Hirose et al. 2011; Gruenheit et al. 2017) and the FLO1 locus in yeast (Smukalla et al. 2008). In these and other microbial examples (Pathak et al. 2013; Heller et al. 2016; Danka et al. 2017), there is genetic linkage between a ligand and a receptor gene, which allows cooperative individuals to aggregate and direct cooperative acts towards each other. Non-carriers of the locus are excluded from the group and thus cannot benefit from cooperation.

Although fascinating in their own right, one aspect of the above examples is that relatedness through common ancestry is not entirely decoupled from high relatedness at the green-beard loci. This is because all study systems involved closely related strains and lineages, sometimes only differing in a single mutation (e.g. FLO1-negative mutants; Smukalla et al. 2008). Hence, it still remains unknown whether the green-beard mechanism would foster cooperation between completely unrelated organisms, where the two relatedness types are not confounded. Here, we study such a scenario where two genetically unrelated, but naturally co-existing, bacterial species *Pseudomonas aeruginosa* (*γ*-proteobacteria) and *Burkholderia cenocepacia* (*β*-proteobacteria) have the potential to cooperate via the sharing of pyochelin, a secreted siderophore used by both species to acquire ferrous iron from iron-limited environments (Weaver and Kolter 2004; Lambiase et al. 2006; Youard et al. 2011; Butt and Thomas 2017). The pyochelin locus has all the characteristics of a green beard, because it consists of the synthesis machinery for the siderophore, a secreted beneficial public good that can be cooperatively shared among individuals in a group and a linked receptor, which is required for the uptake of the pyochelin-iron complex (Ross-Gillespie et al. 2015; Butt and Thomas 2017; Sathe et al. 2019).

Using this system, we first examined whether pyochelin provides fitness benefits in iron-limited medium and can be shared between the members of the different species, thereby constituting a mutually beneficial form of cooperation. Next, we tested whether stable inter-species cooperation arises under iron-limited conditions, where the interests of the two species are aligned through the high relatedness at the green beard locus, which allows to direct pyochelin-mediated cooperation to other carriers of that trait. We then supplemented the medium with iron, which suppresses the production of pyochelin, and thus removes the green-beard effect, such that interactions between the species should be ruled by competition and not cooperation. Finally, we introduce mutants into the system that only make receptors but no pyochelin (i.e. false-beards) and ask whether they can exploit the cooperators of the other species, and thereby bring the green-beard effect to collapse.

## Material and Methods

### Bacterial strains

We used siderophore gene knockout mutants derived from the two laboratory strains *Burkholderia cenocepacia* H111 (LMG23991) and *Pseudomonas aeruginosa* PAO1 (ATCC15692) (Ghysels et al. 2004; Mathew et al. 2014). Both of these species produce pyochelin in addition to their main siderophores (ornibactin for *B. cenocepacia* H111 and pyoverdine for *P. aeruginosa* PAO1) (Leinweber et al. 2017). Since we were solely interested in cooperation mediated via pyochelin, we used strains that were deficient in the production of the primary siderophores, and thus could only produce pyochelin. Specifically, we used: (a) *B. cenocepacia* H111Δ*orbJ*, producing only pyochelin as the ornibactin biosynthesis gene *orbJ* is deleted; (b) *B. cenocepacia* H111Δ*orbJ*Δ*pchAB*, deficient in ornibactin and pyochelin production as the biosynthesis genes *orbJ* and *pchAB* are deleted; (c) *P. aeruginosa* PAO1Δ*pvdD*, producing only pyochelin as the pyoverdine biosynthesis gene *pvdD* is deleted; and (d) *P. aeruginosa* PAO1Δ*pvdD*Δ*pchEF*, deficient in pyoverdine and pyochelin production as the biosynthesis genes *pvdD* and *pchEF* are deleted (Ghysels et al. 2004; Mathew et al. 2014). To be able to distinguish the two species in competition, we used *P. aeruginosa* strains that were chromosomally tagged (single insertion at the attTn7 site) with a constitutive fluorescent mCherry marker (Rezzoagli et al. 2019). All the strains were stored as clonal populations in 25% glycerol stocks at −80°C and freshly revived prior to every experiment.

### Growth measurements

To evaluate the benefit of pyochelin, we compared the growth of bacterial strains in monocultures and in 1:1 inter-species mixed co-cultures both in iron-rich and iron-poor casamino acids (CAA) medium. The CAA was prepared by mixing 5 g/l casamino acids, 1.18 g/l K_2_HPO_4_ × 3H_2_O, 0.25 g/l MgSO_4_ × 7H_2_O and 25 mM HEPES (to buffer pH at neutral levels). We either induced iron-limitation by adding 100 µM 2, 2’-bipyridine (a strong synthetic iron chelator) or made the medium iron rich by adding 100 µM FeCl_3._ Bacterial cells were pre-cultured by inoculating them from glycerol stocks in 50 ml sterile falcon tubes containing 10 ml Lysogeny broth (LB). The tubes were incubated in a 37°C shaken incubator (220 RPM) for approximately 15 hrs. The grown cultures were pelleted using a centrifuge (Eppendorf, 5804 R; 5000 RPM, 22°C, 3 min), washed and re-suspended in 0.8% sterile NaCl solution, and their optical density (measured at 600 nm, OD_600nm_) adjusted to 1. All growth experiments were performed in sterile 96-well plates, where the bacterial strains were inoculated in either 200 µl iron-rich or iron-poor CAA media, at a starting density of OD_600nm_ = 0.01 in four-fold replication for all strains. Plates were incubated statically at 37°C in a microplate reader (SpectraMax Plus 384, Molecular Devices, USA) and OD_600nm_ was monitored every 15 minutes for 24 hrs with 10 sec shaking events prior to each round of growth measurement.

Growth curves were analysed in R using the *grofit* package developed by Kahm *et al.* (2010).We extracted values of growth integrals (i.e. area under the curve) for all treatments and strains. This measure integrates information on the entire growth kinetic, including lag-, exponential and stationary phases, and is ideal to compare growth between strains and species that completely differ in their growth trajectories.

### Supernatant assay to test for inter-species pyochelin cross-use

We first examined whether pyochelin can indeed be cross-used between the two species. To do so, we investigated whether the growth of the siderophore non-producing mutants H111Δ*orbJ*Δ*pchAB* and PAO1Δ*pvdD*Δ*pchEF* can be stimulated in iron-poor medium if supplemented with supernatants containing pyochelin produced by PAO1Δ*pvdD* and H111Δ*orbJ*, respectively. As a control, we provided the siderophore non-producing mutants with their own supernatants. In additional controls, we supplemented the siderophore non-producing mutants with donor supernatants collected (i) from iron-rich medium, where little siderophore is produced, and (ii) from both iron-rich and iron-poor supernatants subsequently replenished with iron to ensure that siderophores are not required for iron acquisition. To generate cell-free supernatants, we grew the donor strains in 50 ml falcon tubes containing either 10 ml iron-rich or iron-poor medium. Tubes were incubated in a shaken incubator (220 RPM) at 37°C for 24 hrs, after which the grown cultures were centrifuged (5000 RPM, 3 min, 22°C) and the supernatants filter-sterilized using 0.2 µm filters. The supernatants were either used immediately or stored at −20°C until use.

For the pyochelin cross-use test, we grew the two recipient strains (PAO1Δ*pvdD*Δ*pchEF* and H111Δ*orbJ*Δ*pchAB*) in medium containing 70% CAA (either iron-rich or iron-poor) supplemented with 30% supernatants from the donor strains. The assays were performed in 96 well plates in 200 µl volumes (medium-supernatant mix) with the recipient strain inoculum being adjusted to OD_600nm_ = 0.01 and in four-fold replication for all combinations. Growth of the recipient strains was recorded in a plate reader as described above.

### Competition assays

We co-cultured the pyochelin producing and non-producing strains of *B. cenocepacia* and *P. aeruginosa* in all possible combinations under iron-poor conditions. Specifically, we co-cultured: (a) the two pyochelin producers PAO1Δ*pvdD* and H111Δ*pchAB*, where cooperation through pyochelin sharing is possible; (b) the two pyochelin non-producers PAO1Δ*pvdD*Δ*pchEF* and H111Δ*orbJ*Δ*pchAB*, where cooperative pyochelin exchange is not possible; (c) PAO1Δ*pvdD*Δ*pchEF* and H111Δ*pchAB*, where *P. aeruginosa* can potentially cheat on the pyochelin produced by *B. cenocepacia*; and (d) PAO1Δ*pvdD* and H111Δ*orbJ*Δ*pchAB*, where *B. cenocepacia* can potentially cheat on the pyochelin produced by *P. aeruginosa*. Because the level of pyochelin sharing and cheating can vary in response to strain mixing, we carried out all competition assays under both static and shaken (170 RPM) conditions. All these co-culture experiments were repeated in iron-rich medium, where pyochelin production is repressed and any of the above-mentioned social interaction should not occur. Prior to competition, the cells of the two species were grown as clonal cultures in LB medium for ∼ 15 hrs, washed, re-suspended in 0.8% NaCl and adjusted to OD 600_nm_ = 1. Competitions occurred in sterile flat bottom 24 well plates (Falcon) containing either 1.5 ml iron-rich or iron-poor media. Cells of the two species were mixed at a 1:1 volumetric ratio (OD 600_nm_ = 0.01) and incubated at 37°C for 24 hrs. For every strain combination, sample size was n = 15 for static conditions and n = 10 for shaken conditions.

We used flow cytometry to determine the ratio of cells from the two species before and after competition. Thanks to the chromosomally integrated stable mCherry tags the cells of the *Pseudomonas* strains (PAO1Δ*pvdD*Δ*pchEF-mCherry* and PAO1Δ*pvdD-mCherry*) could unambiguously be distinguished from the cells of *Burkholderia* strains (see (Sathe et al. 2019), for exact descriptions of the protocol and illustrations). We used a LSR II Fortessa flow cytometer (BD Biosciences), where mCherry was excited at 561 nm and fluorescence emission was quantified with a 610/20 nm bandpass filter. We used low thresholds (200) both for the forward and the side scatter and adjusted the voltage of the photomultiplier tube (PMT) to ensure that the entire bacterial population and all mCherry positive cells were recorded.

For every replicate, we recorded 100,000 bacterial events and used those to calculate the initial (prior to competition) and the final proportion (after 24 hrs competition) of the two competing strains. Under static conditions, a certain percent (5-10%) of PAO1 cells did not express mCherry at a detectable level. To avoid overestimations of the untagged *B. cenocepacia* strains in mixtures, we calculated the final frequency of *B. cenocepacia* by subtracting the proportion of mCherry negative cells of *P. aeruginosa* strains in monocultures from the total of mCherry negative cells in mixed cultures. The flow cytometry data were analysed using the software package FlowJo (TreeStar, USA).

### Testing for frequency-dependent fitness patterns

We conducted another set of competition assays, where we varied the starting ratio of the two competing strains from 1:99; 10:90; 50:50; 90:10 to 99:1. This was done because it is well established that social interactions involving public-goods sharing and cheating can underlay negative frequency-dependant selection, where each of the competing strain experiences relative fitness benefits when rare (Gilbert et al. 2007; Ross-Gillespie et al. 2007; Velicer and Vos 2009). For this set of experiments, we followed the exact same procedures as described above. In competitions where *P. aeruginosa* started at a frequency of 99%, it sometimes occurred that flow cytometry data yielded, after the implementation of all corrections, strain frequencies > 100%. For these cases, we assumed that the mCherry positive *Pseudomonas* strain had fixed in the population and the *Burkholderia* strain went extinct, and consequently we set *Pseudomonas* frequency to 100%.

### Statistical analysis

All statistical analyses were performed with R 3.3.2 (http://www.r-project.org). We used linear models for all analyses. When comparing the growth integrals of the different strains in mono- and mixed cultures, we performed an analysis of variance (ANOVA) to test for media (iron-poor vs. iron-rich) and species (*P. aeruginosa* vs. *B. cenocepacia*) effects. We used the Tukey HSD test for post-hoc pairwise comparisons. To test whether pyochelin can be cross-used between the two species, we carried out ANOVAs to compare whether supernatants from donor strains, proficient in producing pyochelin, stimulate growth of the recipient strains more than supernatants from donor strains that are deficient in pyochelin production.

We further performed ANOVAs to examine whether culturing condition (shaken vs. static) and the type of strains in a co-culture (1:1 mix) affect the frequency of *P. aeruginosa* (our focal species) after competition. Note that it is common to calculate a relative fitness index for the focal strain in mixed cultures and use this index for statistical analysis (Ross-Gillespie et al. 2007). However, as *P. aeruginosa* fixed in many populations, relative fitness indices remain undefined and could not be used in our case. That is why we compared the end frequency of *P. aeruginosa* across treatments in our analyses. Moreover, the monoculture growth experiments revealed that *P. aeruginosa* strains grow overall better than *B. cenocepacia* strains. To account for these intrinsic differences in growth rates, we calculated for each strain pair the expected frequency shift assuming no social interactions. *P. aeruginosa* was again taken as the focal species and the expected frequency shift is calculated as *v*_P_end_ = *v*_P_start_*(μ_P_/µ_B_), where *v*_P_end_ and *v*_P_start_ are the end and start frequencies of *P. aeruginosa* in mixed cultures, and *μ*_P_ and *µ*_B_ are the specific growth rates of *P. aeruginosa* and *B. cenocepacia*, respectively. We used spline fits to calculate *µ* from the monoculture growth trajectories (see supplementary Fig. 1). Finally, we used regression models (fitting linear and quadratic terms) to test whether the frequency of *P. aeruginosa* in mixed cultures varies in response to its starting frequency.

## Results

### Pyochelin production is beneficial for growth in iron-limited medium

When grown as monocultures we found that the strains grew significantly better in iron-rich than in iron-poor medium (ANOVA: F_1,28_ = 59.81; *p* < 0.0001), and that the *P. aeruginosa* PAO1 strains grew significantly better than the *B. cenocepacia* H111 strains (ANOVA: F_1,28_ = 23.14; *p* < 0.0001) (Fig. 1 and Supplementary Fig. S1). To test whether pyochelin production influences fitness, we compared the growth of pyochelin producers and non-producers in iron-poor medium and found that the pyochelin producers grew significantly better than the non-producers in both species (*B. cenocepacia*: t_24_ = 7.67, *p* < 0.0001; *P. aeruginosa*: t_24_ = −20.92, *p* < 0.0001; Fig. 1 and Supplementary Fig. S1). While this growth difference disappeared for *B. cenocepacia* in iron-rich medium (t_24_ = −0.32, *p* = 0.9999), it persisted for *P. aeruginosa* in iron-rich medium (t_24_ = −13.67, *p* < 0.0001). Altogether, these results indicate that pyochelin is an important siderophore promoting growth of both *P. aeruginosa* and *B. cenocepacia* in iron-poor medium. Moreover, the ability to produce pyochelin seems also to be beneficial for *P. aeruginosa* PAO1 but not for *B. cenocepacia* in iron-rich medium.

**Figure 1:**
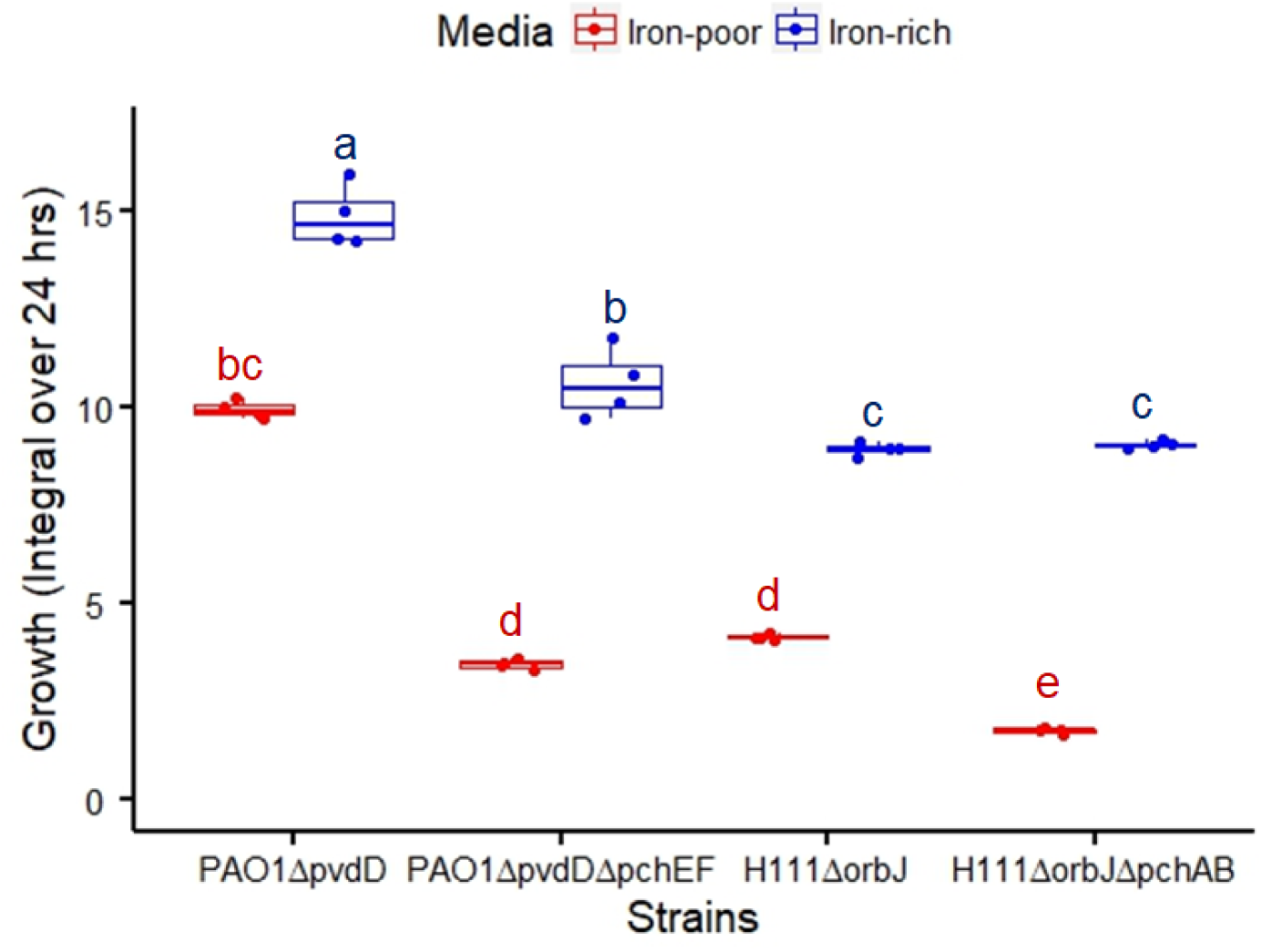
Pyochelin producers grow better than pyochelin non-producers in iron-limited medium. Pyochelin producing strains (PAO1Δ*pvdD*, H111Δ*orbJ*) and pyochelin non-producing strains (PAO1Δ*pvdD*Δ*pchEF*, H111Δ*orbJ*Δ*pchAB*) were grown as monocultures in iron-rich and iron-poor media for 24 hrs and integrals (area under the curve) were used to compare growth performances. Strains grew significantly better in iron-rich compared to iron-poor medium. In iron-poor medium the pyochelin producers of both species grew better than the pyochelin non-producers demonstrating the benefit of pyochelin. The different letters above the boxplots indicate statistically significant growth differences.

### *B. cenocepacia* and *P. aeruginosa* can cross use each other’s pyochelin

Our supernatant assay revealed that pyochelin can be shared between the two species (Fig. 2). Particularly, we observed that in iron-poor medium the siderophore non-producers PAO1Δ*pvdD*Δ*pchEF* (Fig. 2A, F_1,6_ = 1421; *p* < 0.0001) and H111Δ*orbJ*Δ*pchAB* (Fig. 2C, F_1,6_ = 72.39; *p* < 0.0001) were significantly stimulated in their growth when supplemented with supernatants from pyochelin producers of the other species. These comparisons are relative to treatments where the siderophore non-producers received supernatants from hetero-specific donor strains that were themselves deficient for pyochelin production (Fig. 2A+C). In these latter cases, no growth stimulation was observed. When the supernatants from pyochelin producers were collected in iron-rich medium, where no or little pyochelin is produced, growth stimulation was either completely abrogated (Fig. 2B, for PAO1Δ*pvdD*Δ*pchEF*, F_1,6_ = 2.03; *p* = 0.2038) or reduced (Fig. 2D, for H111Δ*orbJ*Δ*pchAB*, F_1,6_ = 2069; *p* < 0.0001). Finally, when supernatants were replenished with iron after their collection, thereby rendering pyochelin superfluous, all growth stimulation effects disappeared (Supplementary Fig. S2). Taken together, these results suggest that pyochelin can be cross-used between the members of the different species and cross-use provides fitness benefits to the receiver. This opens the possibility for inter-species cooperation to arise in iron-limited medium.

**Figure 2:**
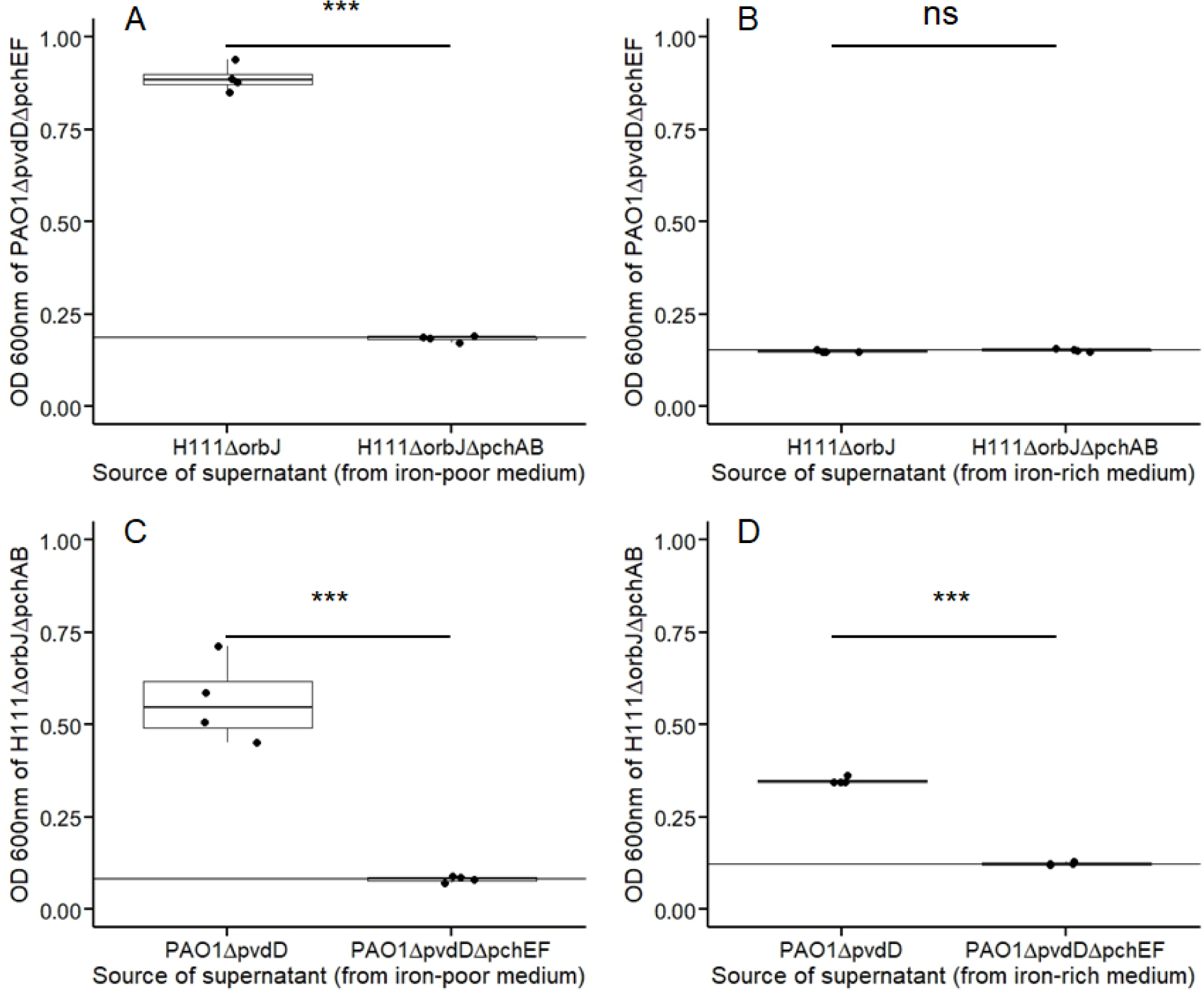
Pyochelin can be shared as public good between strains of *P. aeruginosa* and *B. cenocepacia*. (A) The pyochelin non-producer of *P. aeruginosa* (PAO1Δ*pvdD*Δ*pchEF*) was stimulated in growth when supplemented with pyochelin-containing supernatant from *B. cenocepacia* H111Δ*orbJ*, but not when supplemented with supernatant from *B. cenocepacia* H111Δ*orbJ*Δ*pchAB*, which is deficient for pyochelin production. (B) No growth stimulation was observed when PAO1Δ*pvdD*Δ*pchEF* was supplemented with supernatants from *B. cenocepacia* strains collected from iron-rich medium. (C) Similarly, the pyochelin non-producer of *B. cenocepacia* (H111Δ*orbJ*Δ*pchAB*) was stimulated in growth when supplemented with pyochelin-containing supernatant from *P. aeruginosa* PAO1Δ*pvdD*, but not when supplemented with supernatant from PAO1Δ*pvdD*Δ*pchEF*, which is deficient for pyochelin production. (D) The growth stimulation was reduced or absent when H111Δ*orbJ*Δ*pchAB* was supplemented with supernatants from *P. aeruginosa* strains collected from iron-rich medium.

### *P. aeruginosa* strains consistently outcompete *B. cenocepacia* strains in mixed cultures

In a first experiment, we compared total population growth of co-cultures of *P. aeruginosa* and *B. cenocepacia*. Like for monocultures, we observed that mixed cultures grew significantly better in iron-rich than in iron-poor medium (ANOVA: F_1,24_ = 638.04; *p* < 0.0001; Fig. 3A). In both media, it seemed that the growth patterns were determined by the genetic background of the *P. aeruginosa* strain: cultures containing the pyochelin producer PAO1Δ*pvdD* always grew significantly better than cultures containing the pyochelin non-producer PAO1Δ*pvdD*Δ*pchEF* (ANOVA: F_1,24_ = 495.38; *p* < 0.0001), while the genetic background of *B. cenocepacia* played no role (ANOVA: F_1,24_ = 1.25; *p* = 0.2728). This pattern yielded a first hint that competitions might be determined by *P. aeruginosa*.

**Figure 3:**
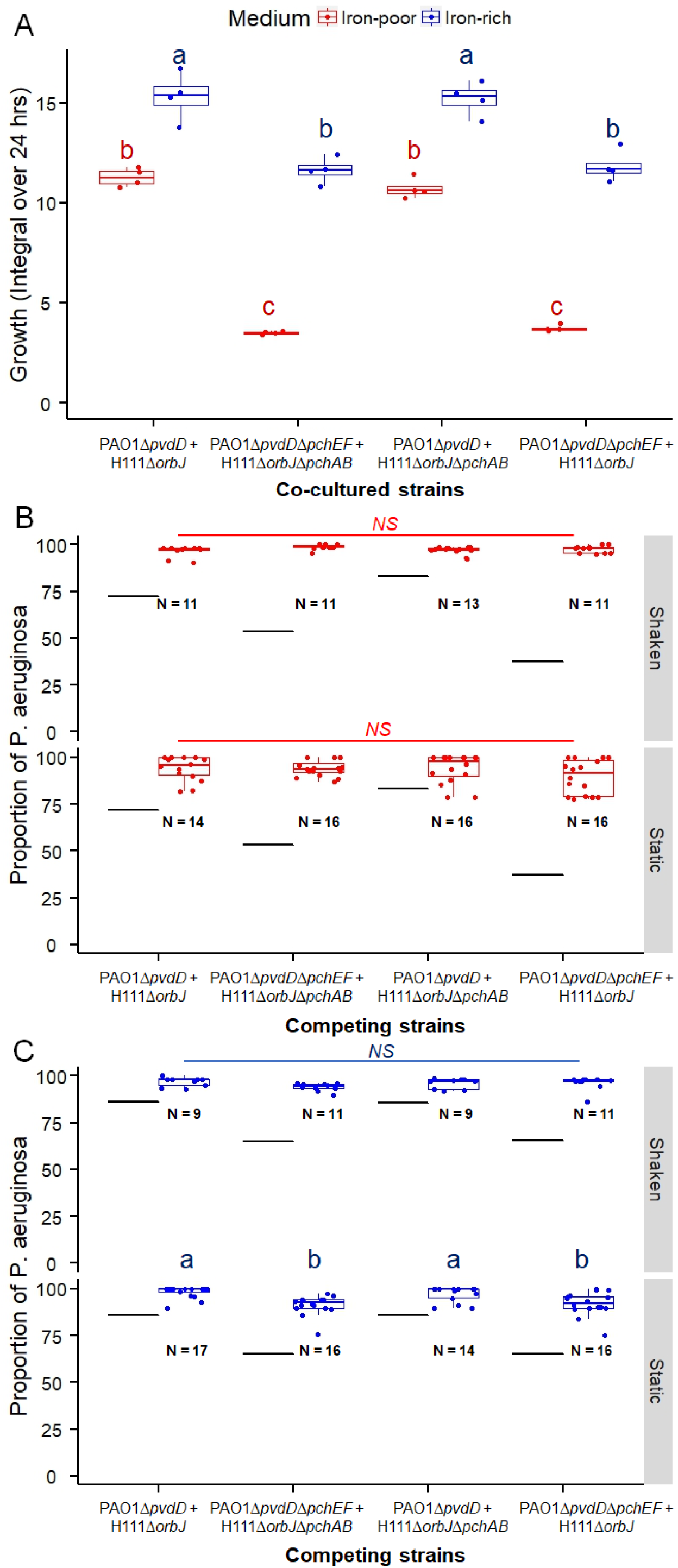
Total population growth and proportions of *P. aeruginosa* strains in co-cultures with *B. cenocepacia* strains. To examine the role of pyochelin in species interactions, we mixed the two species in four different combinations at equal proportions. The four mixes included: (i) PAO1Δ*pvdD* vs. H111Δ*orbJ*, pyochelin-mediated green beard cooperation is possible; (ii) PAO1Δ*pvdD*Δ*pchEF* vs. H111Δ*orbJ*Δ*pchAB*, pyochelin-mediated green beard cooperation is disabled; (iii) PAO1Δ*pvdD* vs. H111Δ*orbJ*Δ*pchAB*, the latter strain can cheat on the pyochelin produced by the former; (iv) PAO1Δ*pvdD*Δ*pchEF* vs. H111Δ*orbJ*, the former strain can cheat on the pyochelin produced by the latter. (A) Mixed cultures grew significantly better in iron-rich than in iron-poor medium. In both media, the population growth was highest in mixes involving the *P. aeruginosa* pyochelin producer (PAO1Δ*pvdD*). (B) Under iron-poor conditions, *P. aeruginosa* strains significantly outcompeted *B. cenocepacia* strains both under shaken and static culturing conditions. There were no differences between strain mixes, indicating that pyochelin-mediated interactions play no significant role. (C) Under iron-rich conditions, *P. aeruginosa* strains significantly outcompeted *B. cenocepacia* strains both under shaken and static culturing conditions. Black lines indicates the expected proportions of *P. aeruginosa* strains after 24 hrs based on the growth rate differences observed in monocultures. NS = not significant. Letters above boxplots indicate significant differences between strain type mixes.

We then compared the proportions of the co-cultured strains before and after 24 hrs of competition. We found that the *P. aeruginosa* strains consistently and strongly outcompeted the *Burkholderia* strains in both iron-poor and iron-rich medium (comparison across all strain combinations relative to the expected frequency shifts based on monoculture growth; ANOVA for iron-poor medium: F_1,113_ = 145.80; *p* < 0.0001, Fig. 3B; ANOVA for iron-rich medium: F_1,108_ = 96.20; *p <* 0.0001, Fig. 3C). Under iron-limited conditions, the dominance of *P. aeruginosa* did not differ between the four different co-culture types (ANOVA: F_3,103_ = 1.29; *p =* 0.2794), but was more pronounced in shaken than in static cultures (ANOVA: F_1,105_ = 16.87; P < 0.0001). Overall, these results indicate that the competitive interactions between the two species subdue any fitness effects that might arise from social interactions mediated by either pyochelin cooperation (PAO1Δ*pvdD* vs. H111Δ*orbJ*) or cheating (PAO1Δ*pvdD* vs. H111Δ*orbJ*Δ*pchAB* and PAO1Δ*pvdD*Δ*pchEF* vs. H111Δ*orbJ*).

### *P. aeruginosa* outcompetes *B. cenocepacia* across a wide range of starting frequencies

Given that *P. aeruginosa* was least dominant in iron-limited medium under static culturing conditions, we wondered whether co-existence of the two species could arise at specific strain ratios and whether social interactions could be involved in reaching an equilibrium frequency at which the species can co-exist. We therefore mixed the strains of the two species across a range of starting frequencies (1:99, 10:90, 50:50, 90:10, 99:1) and monitored their proportions after 24 hrs of growth in iron-rich and iron-poor static media. We found that frequency-dependent fitness patterns were driven by the identity of the *P. aeruginosa* strain and not by the type of social interactions involved (Fig. 4). Specifically, we observed positive relationships between initial and final *P. aeruginosa* frequencies when PAO1Δ*pvdD* was the competitor (Fig. 4A, versus H111Δ*orbJ*: F_1,43_ = 59.04; *p* < 0.0001; Fig. 4C, versus H111Δ*orbJ*Δ*pchAB*: F_1,45_ = 50.29; *p* < 0. 0001). The fitness trajectories indicate that PAO1Δ*pvdD* drives *B. cenocepacia* strains to extinction. In contrast, no frequency-dependent patterns were observed when PAO1Δ*pvdD*Δ*pchEF* was the competitor (Fig. 4B, versus H111Δ*orbJ*Δ*pchAB*: F_1,45_ = 0.32; *p* = 0.5708; Fig. 4D, versus H111Δ*orbJ*: F_1,45_ = 0.24; *p* = 0. 6240). In these cases, PAO1Δ*pvdD*Δ*pchEF* reaches similarly high frequencies (91.86±0.82; mean±SE) regardless of its starting frequencies, but rarely drives *B. cenocepacia* strains to extinction. Finally, in iron-rich static control cultures where siderophores are not required, we detected positive relationships between initial and final *P. aeruginosa* frequencies for all four mixing combinations with the data suggesting that PAO1Δ*pvdD* drives *B. cenocepacia* strains to extinction (Fig. 4E, PAO1Δ*pvdD* vs. H111Δ*orbJ*: F_1,45_ = 30.83, *p* < 0.0001; Fig. 4F, PAO1Δ*pvdD*Δ*pchEF* vs. H111Δ*orbJ*Δ*pchAB*: F_1,43_ = 55.74, *p* < 0.0001; Fig. 4G, PAO1Δ*pvdD* vs. H111Δ*orbJ*Δ*pchAB*: F_1,41_ = 112.8, *p* < 0.0001; Fig. 4H, PAO1Δ*pvdD*Δ*pchEF* vs. H111Δ*orbJ*: F_1,41_ = 40.60; *p* < 0.0001).

**Figure 4:**
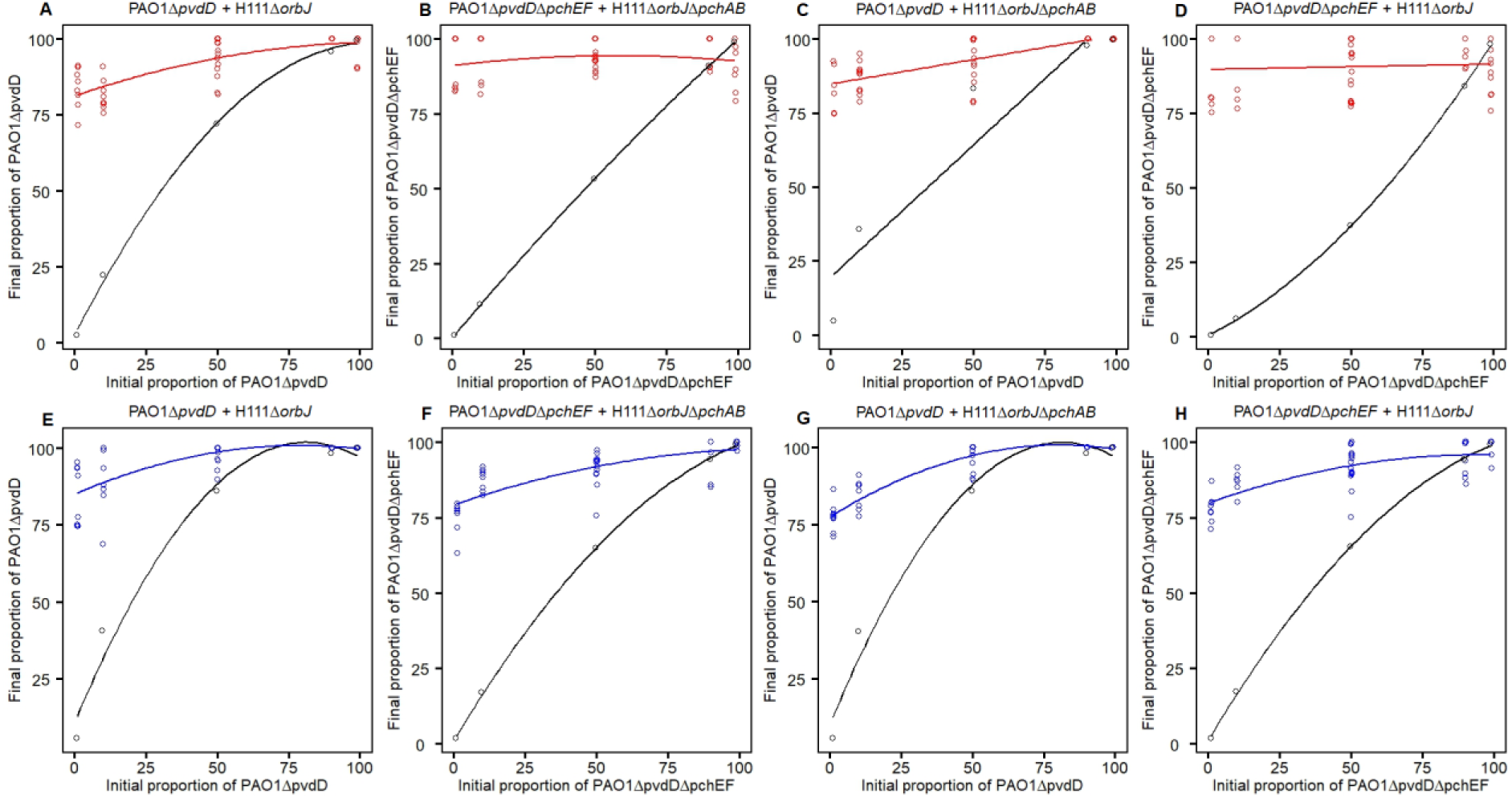
The effect of varying starting frequencies on the outcome of inter-species interactions. To test whether *P. aeruginosa* strains dominate at all strain frequencies, we repeated the competition assays by mixing strains of the two species at different initial starting ratios (1:99, 10:90, 50:50, 90:10 and 99:1) under iron-poor (A-D) and iron-rich (E-H) conditions. (A + C) The final proportions of *P. aeruginosa* pyochelin producers (PAO1Δ*pvdD*) depended on their initial strain frequencies, and *B. cenocepacia* strains are finally driven to extinction. (B + D) The final proportions of *P. aeruginosa* pyochelin non-producers (PAO1Δ*pvdD*Δ*pchEF*) did not depend on their initial strain frequencies, and *B. cenocepacia* strains were maintained at low frequencies (∼8%) in the populations. (E-H) The final proportions of both *P. aeruginosa* strains depended on their initial strain frequencies, and in all cases they seem to drive *B. cenocepacia* strains to extinction. Black open circles and solid lines represent the expected shifts in *P. aeruginosa* proportions based on the growth rate differences between the strains in monocultures.

## Discussion

We set out to test whether green-beard cooperation can occur between two phylogenetically unrelated species. We used the production and sharing of the siderophore pyochelin between the two bacterial species *P. aeruginosa* PAO1 and *B. cenocepacia* H111 as a model system. The pyochelin locus has all the features of a green-beard locus because it entails (i) a cooperative trait, pyochelin; (ii) a green beard, the pyochelin receptor; and (iii) a regulatory and physical link between the two traits as they are part of the same regulon and locus, respectively (Youard et al. 2011; Butt and Thomas 2017). We thus presumed that the two species could derive equal benefits from the shared pool of pyochelin under iron-limited conditions, where pyochelin is important for growth, and thereby maximize population growth and eventually allow species to settle at a stable frequency. While we found that pyochelin indeed increases population growth and is shared as a public good between members of the two species, there was no evidence that pyochelin-mediated green-beard cooperation played a relevant role in determining the fate of our two-species communities. In fact, we observed that *P. aeruginosa* was the dominant species driving *B. cenocepacia* to extinction under most conditions. The dominance of *P. aeruginosa* was beyond the level one would expect based on growth rate differences alone. This indicates that *P. aeruginosa* can sense the presence of *B. cenocepacia* (Cornforth and Foster 2013; Leinweber et al. 2018) and mount competitive responses that subdue any mutual benefits that could arise from green-beard cooperation. Our results highlight that selection for competitive traits might outrace selection for cooperation in inter-species interactions (Foster and Bell 2012; Oliveira et al. 2014), even in the case where the two competing species share the same cooperative trait.

Green beard cooperation was proposed to be rare because it can create inter-genomic conflicts (Dawkins 1976; Burt and Trivers 2006; Gardner and West 2010), although the strength of the conflict seems to depend on biological details including the level of genealogical relatedness (Biernaskie et al. 2011). While our experimental system does not allow to directly test the inter-genomic conflict theory, our results nonetheless highlight that there seem to be divergent interests between the pyochelin green beard locus and the rest of the genome. With respect to the pyochelin locus, we found that the secreted siderophore can be shared and provide fitness benefits across the species boundaries (Figs. 1 + 2). Thus, cooperation between the species could be beneficial. On the other hand, our fitness data indicate that *P. aeruginosa* triggered a specific competitive response in co-culture with *B. cenocepacia*, which enabled *P. aeruginosa* to wipe out its competitor under most conditions (Fig. 4). Our assertion is based on the observation that the dominance of *P. aeruginosa* cannot be explained by growth rate differences alone (Fig. 3 + Fig. S1). In fact, for the strain pairings PAO1Δ*pvdD*Δ*pchEF* vs. H111Δ*orbJ*Δ*pchAB* and PAO1Δ*pvdD*Δ*pchEF* vs. H111Δ*orbJ* one would even expect stable species co-existence or mild *B. cenocepacia* dominance in the absence of interference competition. While we can only speculate about the specific strategies deployed by *P. aeruginosa*, it is well known that this species possesses several mechanisms to engage in interference competition, including the secretion of diffusible toxins such as cyanide (Bernier et al. 2016) and pyocyanin (O’Brien and Fothergill 2017) and the use of contact-dependent killing systems like the type VI secretion system (Basler et al. 2013). Overall, these considerations suggest that selection pressures for sensing competitors, as opposed to sensing co-operators, were much stronger in the evolutionary past of *P. aeruginosa*, with the competitive responses triggered simply overruling any benefits that could accrue through pyochelin green-beard cooperation. In this context, it would be interesting to see whether the benefits of pyochelin green-beard cooperation could be recovered in engineered strains deficient for interference competition.

Another reason for why green-beard cooperation was proposed to be rare is because false beards (i.e. cheaters) can evolve that exploit the cooperative act without contributing to it (Gardner and West 2010). We also examined this possibility by competing pyochelin producers of both species against the respective inter-specific non-producer. Although exploitation of pyochelin produced by the other species has likely occurred (Fig. 2), we did not detect the expected fitness effects that characterize successful cheating (West et al. 2006; Ghoul et al. 2014). Particularly, cheating is expected to lower population productivity, while increasing the relative fitness advantage of cheaters. While previous work showed that these fitness effects manifest in intra-specific interactions between pyochelin producers and non-producers (Ross-Gillespie et al. 2015; Sathe et al. 2019), they did not arise in the inter-specific interactions examined here (Fig. 3). Instead, we observed that *P. aeruginosa* outcompeted *B. cenocepacia* under all conditions, with the pyochelin producer PAO1Δ*pvdD* being the stronger competitor than the pyochelin non-producer PAO1Δ*pvdD*Δ*pchEF* (Fig. 4). Again, these findings indicate that pyochelin-mediated social interactions, including cheating, were subdued by interference competition mediated by other loci in the genome.

Our results further suggest that pyochelin regulation differs between the species, which could also explain why green-beard cooperation was not efficient in our system. *P. aeruginosa* and *B. cenocepacia* belong to phylogenetically different groups of proteobacteria, yet they share the pyochelin locus with identical genetic structure (Youard et al. 2011; Butt and Thomas 2017): the locus consists of two biosynthesis operons *pchABCD* and *pchEFG*, the export/import operon *fptABCX* and the regulator *pchR*. It seems fair to assume that trait co-occurrence is explained by horizontal gene transfer, which is common in bacteria including the transfer of social traits (Nogueira et al. 2009; Mitri and Foster 2013; Soucy et al. 2015). However, despite the genetic architecture of the locus being conserved we argue that natural selection might have acted at the regulatory level, following horizontal gene transfer. Regulatory modifications can differentially affect the costs and benefits of pyochelin production in the two species. For example, one intriguing feature is that pyochelin is considered a secondary siderophore (Cornelis 2010), with its expression being suppressed by the respective primary siderophore (pyoverdine and ornibactin) (Dumas et al. 2013; Tyrrell et al. 2015). However, this suppression seems to be stronger in *P. aeruginosa* (Dumas et al. 2013) than in *B. cenocepacia* (Sathe et al. 2019). Moreover, the results from our study reveal that the two species respond differently to high iron availability: *P. aeruginosa* still produced a certain amount of pyochelin, whereas *B. cenocepacia* stalled production completely (Fig. 2). Our findings further indicate that the net benefit of pyochelin production is larger for *P. aeruginosa* than for *B. cenocepacia* (Fig. 1). Additional differences could include variation in pyochelin molecule and receptor production rates. For example, a lower rate of receptor synthesis in the *P. aeruginosa* pyochelin non-producer PAO1Δ*pvdD*Δ*pchEF* compared to *B. cenocepacia* strains could explain why this *Pseudomonas* strain cannot efficiently cheat and why *B. cenocepacia* strains can be maintained in the population at low frequency (Fig. 4B+D). Altogether, these considerations highlight that differential adaptations at the gene regulation level could lead to customized trait expression patterns for the individual species, which could in turn jeopardize the possibility for green-beard cooperation between species.

In conclusion, we examined an extreme form of the green-beard mechanism, where two unrelated bacterial species share high relatedness at the cooperative locus (pyochelin), but are largely unrelated throughout the rest of their genomes. We found no evidence that this form of green-beard cooperation can be evolutionary stable. Our analyses indicate that selection for competitive traits in the rest of the genome and regulatory co-option of the green-beard locus together overrule any benefits that bacteria could derive from between-species cooperation.

## Figure and Figure legends

**Supplementary Figure 1:**
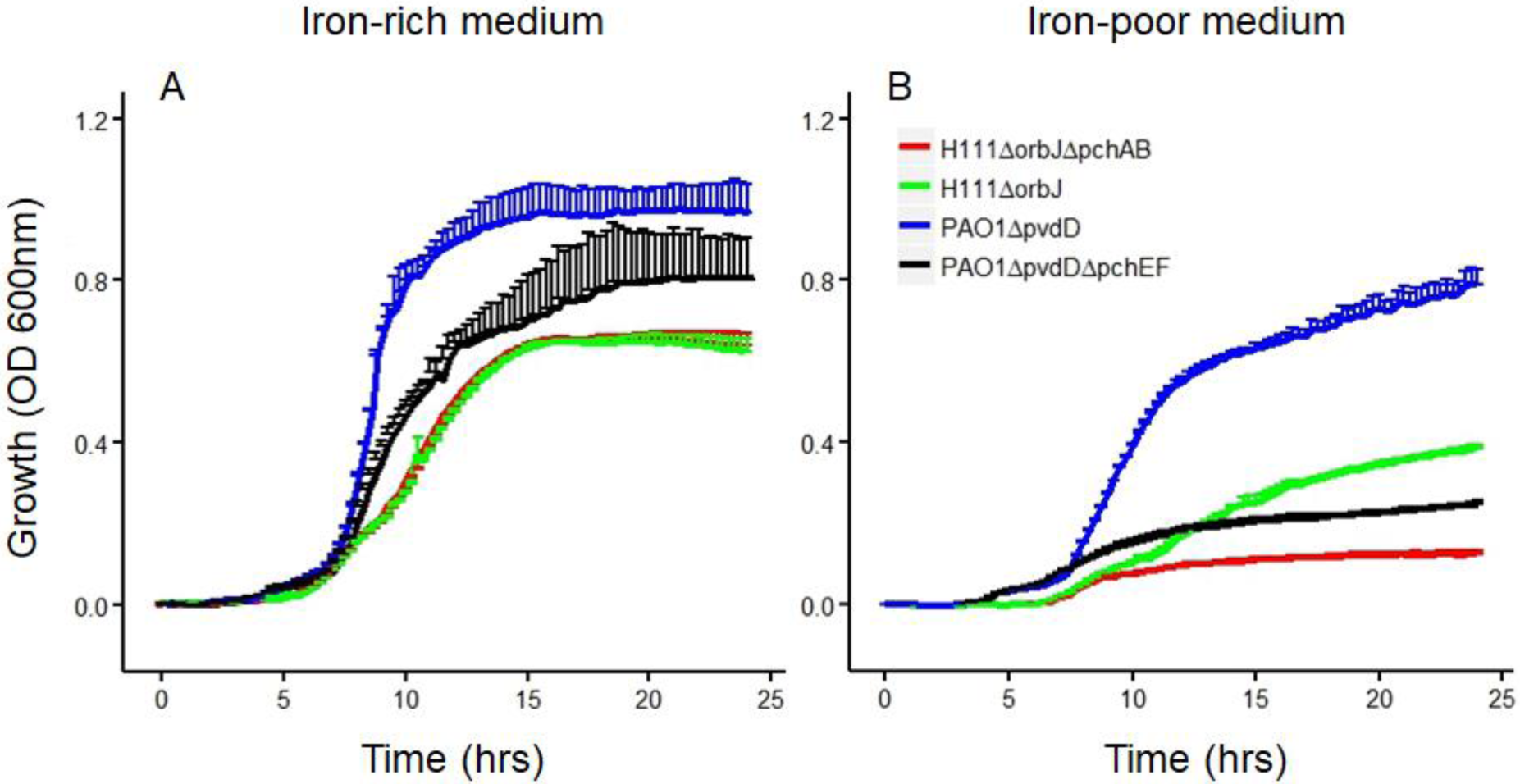
Growth pattern of pyochelin producers and non-producers. We compared the growth of strains producing either pyochelin (PAO1Δ*pvdD*, H111Δ*orbJ*) or no pyochelin (PAO1Δ*pvdD*Δ*pchEF*, H111Δ*orbJ*Δ*pchAB*) as monocultures in iron-rich (A) and iron-poor (B) media. Growth was measured as the optical density (OD) at 600 nm every 15 minutes for 24 hrs. The data is shown as means ± standard deviations across 4 replicates.

**Supplementary Figure 2:**
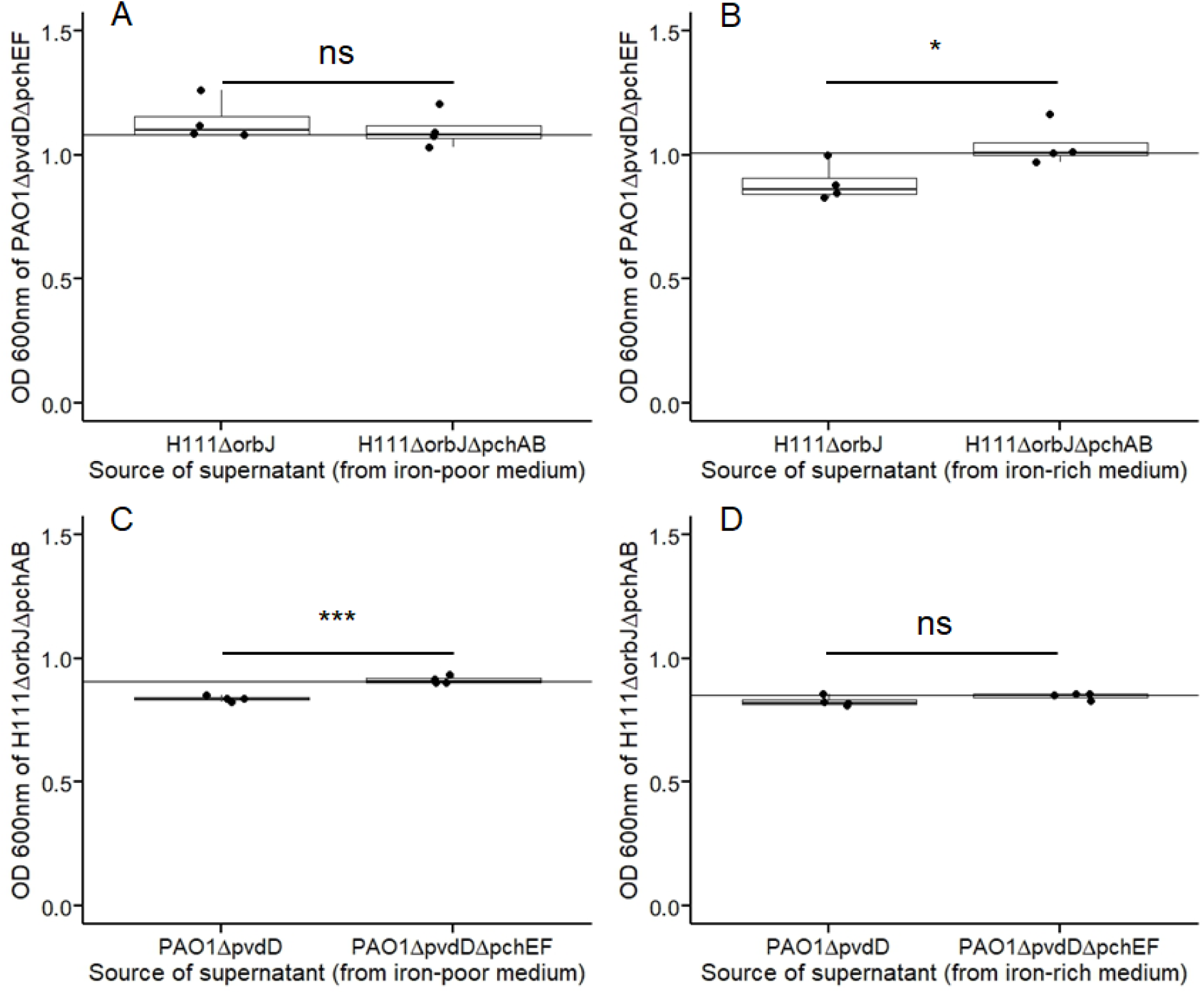
In iron-rich medium where siderophores are redundant the growth stimulatory effects of pyochelin disappear. (A) The pyochelin non-producer of *P. aeruginosa* (PAO1Δ*pvdD*Δ*pchEF*) grew equally well in iron-rich medium regardless of whether pyochelin from *B. cenocepacia* was present (H111Δ*orbJ*) or absent (H111Δ*orbJ*Δ*pchAB*) in the supernatant (F_1,6_ = 0.37; *p* = 0.5625). (B) The pyochelin non-producer of *P. aeruginosa* (PAO1Δ*pvdD*Δ*pchEF*) grew slightly worse in iron-rich medium when supplemented with iron-rich supernatants from H111Δ*orbJ* compared to H111Δ*orbJ*Δ*pchAB* (F_1,6_ = 7.06; *p* = 0.0376). (C) The pyochelin non-producer of *B. cenocepacia* (*H111*Δ*orbJ*Δpc*hAB*) grew slightly worse in iron-rich medium when pyochelin of *P. aeruginosa* (PAO1Δ*pvdD*) was present as opposed to when it was absent (PAO1Δ*pvdD*Δ*pchEF*) in the supernatant (F_1,6_ = 59.33; *p* = 0.0003). (D) The pyochelin non-producer of *B. cenocepacia* (*H111*Δ*orbJ*Δpc*hAB*) grew equally well in iron-rich medium regardless of whether the iron-rich supernatant originated from PAO1Δ*pvdD* or PAO1Δ*pvdD*Δ*pchEF* (F_1,6_ = 2.65; *p* = 0.1543).

